# Chronic Stress Exacerbates Long-term Microvascular Network Dysfunction Following Brain Trauma

**DOI:** 10.64898/2026.06.05.730535

**Authors:** Matthew W Rozak, Adrienne Dorr, Shruti Patel, Margaret Koletar, Ahmadreza Attarpour, Yunze Du, Jonas Osman, Mary Hill, James R Mester, Matthew J. Burke, Clement Hamani, John G Sled, Maged Goubran, Bojana Stefanovic

## Abstract

**Background:** Preexisting factors are among the strongest predictors of recovery following traumatic brain injury (TBI), with chronic stress closely associated with permanent disability and worse long-term outcomes. While chronic cerebrovascular dysfunction is linked to these poor trajectories, as impaired blood flow regulation drives secondary disease progression, the mechanisms regulating this vascular failure remain incompletely understood. Crucially, how premorbid chronic stress and TBI disrupt the fundamental coordination of the cerebrovascular network post-injury remains unknown.

**Methods:** To elucidate this coordination at rest and in response to increased neuronal activity, we used a model of three repeated moderate closed cortical impacts comorbid with chronic stress induced by social isolation (SI) post-weaning. Our previously developed vascular analysis pipeline (NOVAS3D) was employed to estimate changes in vascular radii across cerebrovasculature proximal to neuronal activation. Arteries and veins were annotated in the imaged volumes to allow for blood flow simulations.

**Results:** Using graph-based network analysis, we demonstrate that TBI, when compounded with chronic stress, critically disrupts the long-range coordination of the capillary network. Specifically, the functional coordination of radius changes between nearby, non-adjacent capillaries was reduced by 40±20% in TBI+SI mice relative to controls. Consequently, simulations estimated that the vascular networks in TBI+SI mice experienced a 68±7% reduction in arterial red blood cell velocity (VRBC) responses to neuronal activation. These network-wide impairments were fundamentally driven by severely blunted vessel reactivity, including a 28±7% decrease in the magnitude of arteriolar dilations and a 47±7% decrease in the magnitude of arteriolar constrictions.

**Conclusion:** These findings provide a mechanistic foundation for worse clinical outcomes seen in TBI patients with comorbid chronic stress, identifying arteriolar reactivity and long-range capillary coordination as critical therapeutic targets to mitigate secondary injury and improve long-term recovery.

## Introduction

TBI is a leading cause of disability, with an estimated 64-74 million injuries each year globally, of which roughly 21 million receive moderate to severe injuries, and 37.9 million globally live with permanent disability caused by TBI ^1–6^. Repeated injuries further increase the probability that an individual will experience long-term deficits and disability ^7^. In moderate TBI (Glasgow coma scale 9-12), around 20% of patients move to long-term care homes, and 44% are unable to resume working ^8–10^. Patients with moderate TBI also have an increased risk of developing dementia (1.81 fold), Parkinson’s disease (1.48 fold), and other neurodegenerative diseases later in life ^11–14^. Post-concussive cerebrovascular dysfunction has emerged as an imaging biomarker that can distinguish between individuals who recover well and those who do not ^11,15–17^.

In the acute stage, TBI leads to haemorrhage, edema, vascular loss, breakdown of the blood-brain barrier, and alterations in pericyte density ^15,18–20^. In the chronic stage, as swelling and acute responses subside, angiogenesis and support cells regeneration take place, with an altered structure of new vasculature resulting in blood flow changes ^15,21–23^. Vascular abnormalities commonly include vascular density alterations, atrophy, modulations in functional hyperaemia, and are correlated with axonal disruptions ^24,25^; however, vascular structural alterations do not correlate with blood flow alterations ^26,27^. Residual blood flow and perfusion alterations during functional hyperaemia, particularly when metabolic supply does not meet the needs of neurons, can lead to localized hypoxia, further damaging tissue in the chronic stage of the disease ^15,18,20,21,28,29^. Elucidating vascular network remodelling patterns and flow distribution patterns may provide targets for preventing secondary injury.

The likelihood of sustained deficits post-TBI is increased in the presence of baseline comorbidities. Chronic stress, such as that induced by social isolation (SI), can increase the vulnerability of brain tissue to long-term damage after TBI, through heightened baseline susceptibility to neuroinflammation and stress-induced vascular changes ^30–32^. SI is a prevalent cause of chronic stress, especially in domestic violence victims and older populations who also experience increased risk of TBI ^33–36^. At baseline, SI induces a chronic pro-inflammatory state that primes the brain for a more severe secondary injury ^30–32^. Preclinical models of SI via single housing post-weaning have demonstrated an increase in inflammatory factors, including tumour necrosis factor-alpha and Toll-like receptors interleukin 6 ^37^. If the chronic stress is high enough, the brain can be persistently inflamed ^38^. Socially isolated rodents also display elevated baseline levels of corticosterone, which can attenuate vascular reactivity and promote atherosclerosis^31,39–41^. While previous research has established that chronic developmental stress exacerbates neurobehavioural deficits and glial-mediated neuroinflammation following TBI in adulthood, its compounding impact on neurovascular coupling and microvascular coordination remains entirely unknown. To address this gap, we investigated cerebrovascular network coordination in a mouse model of repeated TBI comorbid with chronic stress induced by SI ^42^. Such inflammatory susceptibility and stress-induced vascular changes motivate the examination of the combined effect of SI and TBI on vascular function.

Varying data have been reported regarding vascular function alterations in TBI, with some studies observing arteriolar and capillary dysfunction ^43^, while others find venular dysfunction ^23^. Despite extensive characterization of individual vascular responses and their dysfunction, how these mechanisms integrate to redistribute blood flow in both health and disease remains unclear. Traditional methods relying on spatially averaged signals capture the broad hypoperfusion characteristic of chronic TBI ^21,22^, but effectively flatten the data, completely filtering out the crucial, vessel-to-vessel spatial dynamics, such as proximal dilations and distal constrictions, that govern flow redistribution. To gain a comprehensive understanding of the microvascular functional deficits and identify targets for treatment, a detailed network-level approach is essential.

To address the lack of single-vessel spatial resolution in prior studies and elucidate how these mechanisms integrate, we set out to interrogate cerebrovascular network coordination in a mouse model of repeated TBI comorbid with chronic stress. We employed volumetric two-photon fluorescence microscopy (2PFM) to map morphological changes in the cerebrovascular network, following increases in local neuronal activity via opsin photoactivation or forepaw stimulation. Cerebrovascular structural and functional changes were tracked on a graph representation of the vasculature, allowing an unprecedentedly detailed analysis of the neurovascular coupling and elucidating mechanisms underlying cerebrovascular supply modulation in the chronic stages of TBI comorbid with chronic stress. Arteries, capillaries, and veins were classified to allow for tracking of changes across branch orders within the network and enable blood flow simulations that link upstream and downstream changes in vessels. Our network-level analysis revealed that mice with TBI comorbid with chronic stress exhibit reduced arteriolar reactivity and a decreased capacity to regulate blood flow from arteries.

Furthermore, we found that long-distance coordination across the capillary network is significantly impaired, suggesting a fundamental breakdown in the spatial organization of blood flow redistribution following combined injury and stress.

## Results

To evaluate microvascular coordination across the experimental cohorts, the NOVAS3D pipeline was first employed to extract, classify, and spatially map the cerebrovascular topologies from raw in vivo two-photon imaging volumes ^44^. Visualizing these structurally distinct networks alongside their localized functional deviations (**Figure 1**) provides the foundational framework necessary for assessing the specific microvascular impairments detailed below.

**Figure 1:**
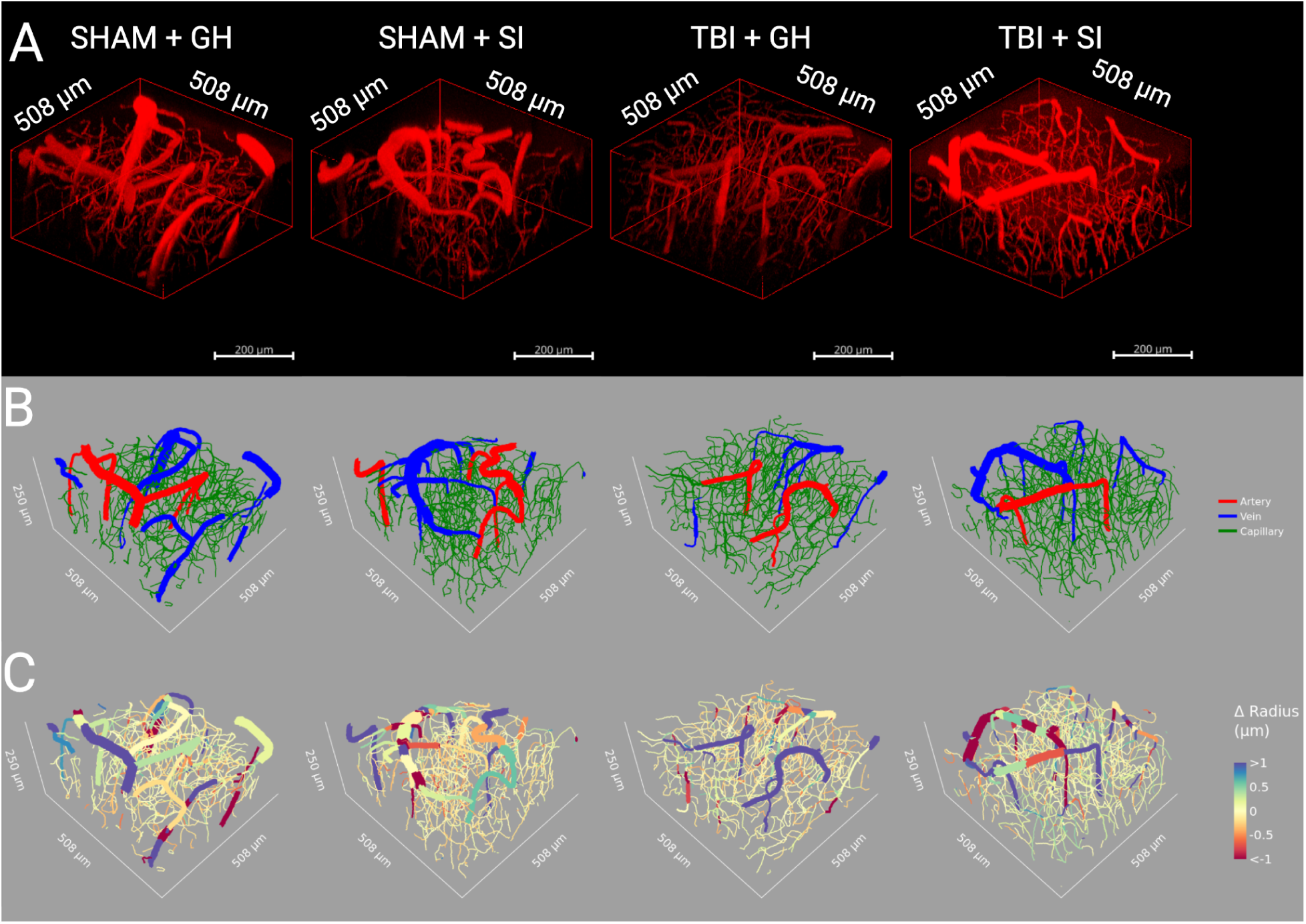
Topological classification and spatial mapping of microvascular reactivity across chronic stress and TBI cohorts. **A.** Representative in vivo 2PFM volumetric acquisitions of the cerebrovascular network within the primary somatosensory cortex. Volumes are shown for each of the four experimental groups: SHAM Group Housed (GH), SHAM SI, TBI GH, and TBI SI. **B.** Corresponding 3D spatial graphs extracted using the NOVAS3D pipeline. Individual vessel segments are classified and colour-coded to delineate the vascular hierarchy: arteries (red), veins (blue), and capillaries (green). **C.** Network-wide spatial heatmaps visualizing the absolute change in vessel radius (Δ Radius) mapped directly onto the extracted topologies, illustrating the localized morphological and functional deviations within the microvascular beds of each cohort.

### Characterization of Vascular Network Function in Control Conditions

To characterize the healthy neurovascular function, we first evaluated the cortical vascular network in SHAM Group Housed (GH) mice (**Figure 2**). We generated 3D reconstructions of the vascular networks, differentiating between arteries, capillaries, and veins (**Figure 2A**), and mapped the spatial distribution of evoked radius changes and simulated VRBC changes following 458-nm (4.3 mW/mm^2^) optogenetic stimulation (**Figure 2B-C**). This analysis established the baseline magnitudes of vascular dilations, constrictions, and velocity responses stratified by branch order (**Figure 2D-I**) in the healthy cortex.

**Figure 2.**
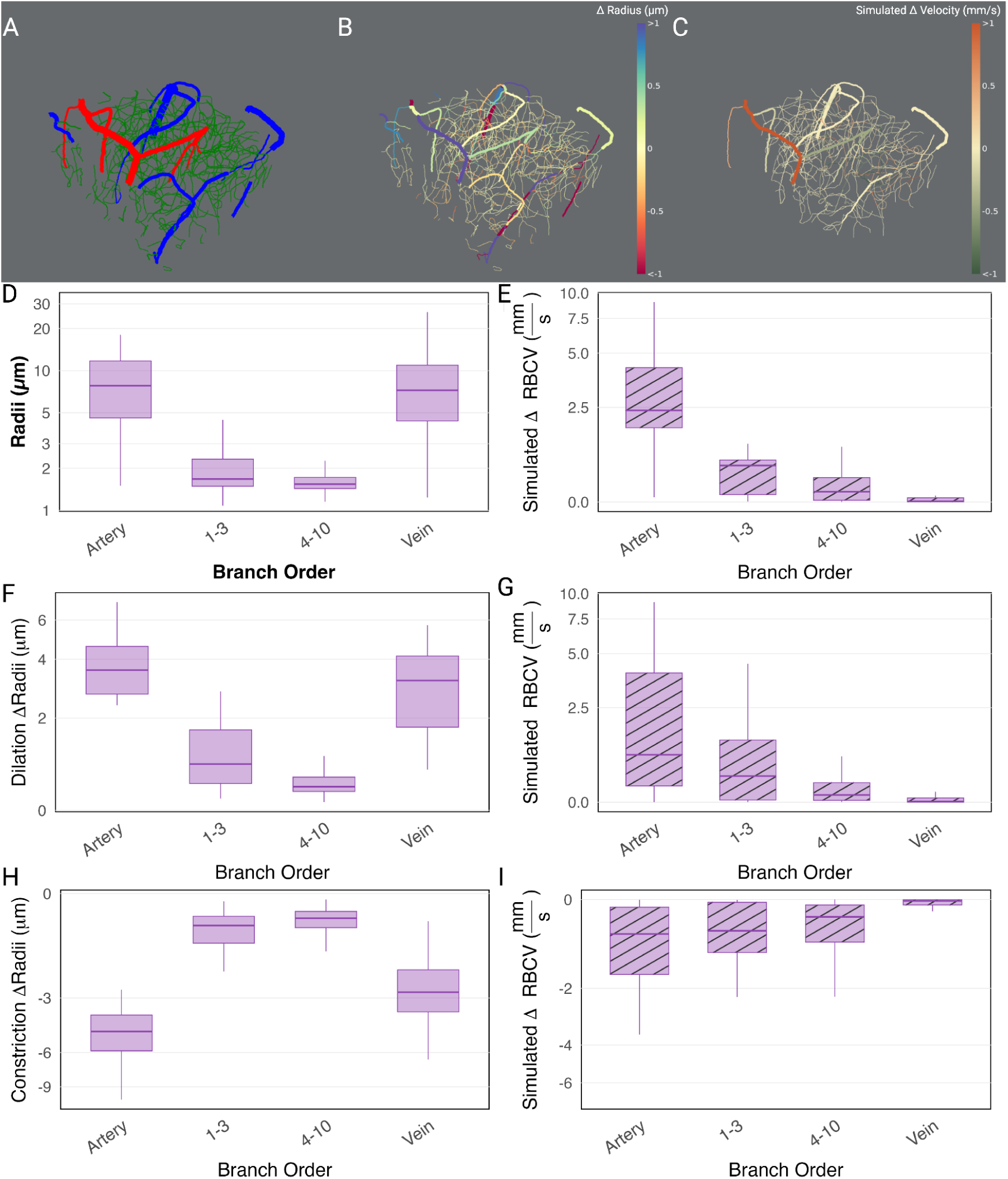
Vascular Network Function in SHAM Group Housed (GH) mice. **A.** Representative 3D reconstruction of the cortical vascular network in a SHAM GH mouse. Vessels are colour-coded by type: arteries (red), capillaries (green), and veins (blue). **B.** Spatial map of evoked radius changes and **C.** simulated VRBC changes following 4.3 mW/mm^2^ optogenetic stimulation (458 nm). **D.** Boxplots of baseline radius estimates stratified by branch order. **F,H.** Magnitude of significant dilations (**F)** and constrictions (**H)** in vascular radius. **E.** Simulated baseline VRBC stratified by branch order. **G, I.** Magnitude of significant increases (**G)** and decreases (**I)** in simulated VRBC following stimulation.

We further examined the coordination of these vascular responses using network analysis (**Figure 3**). Topological representations revealed how vessels, depicted as nodes connected by edges (**Figure 3B**), coordinated their radii changes. Vascular response assortativity (correlation of radius changes between vessel segments) varied as a function of both network distance (degree of separation, **Figure 3C**) and Euclidean distance (**Figure 3D**). In both cases, assortativity decreased with distance. To robustly capture spatial dynamics, vertex-wise data of significant vascular responses (>2x baseline standard deviation(SD)) were spatially downsampled and analyzed using a Generalized Additive Model (GAM). This approach demonstrated a clear spatial gradient of response in the healthy condition, with significant dilations dominating in the central activation core and transitioning to significant constrictions in the periphery **(Figure 3E-F)**. Specifically, while the central activation core demonstrates variable expansive responses, the 95% confidence intervals reveal a robust, significant peripheral constriction beginning at 106.9 μm and extending to the outer bounds of the modelled region (359 μm). Beyond functional metrics, we assessed general vascular network structure. Despite the functional diversity, we found no significant variance in vascular density, length density, or vessel number density across any of the experimental groups (**Figure S2 A-C**), indicating that the gross topology of the network remains robust even in pathological states.

**Figure 3.**
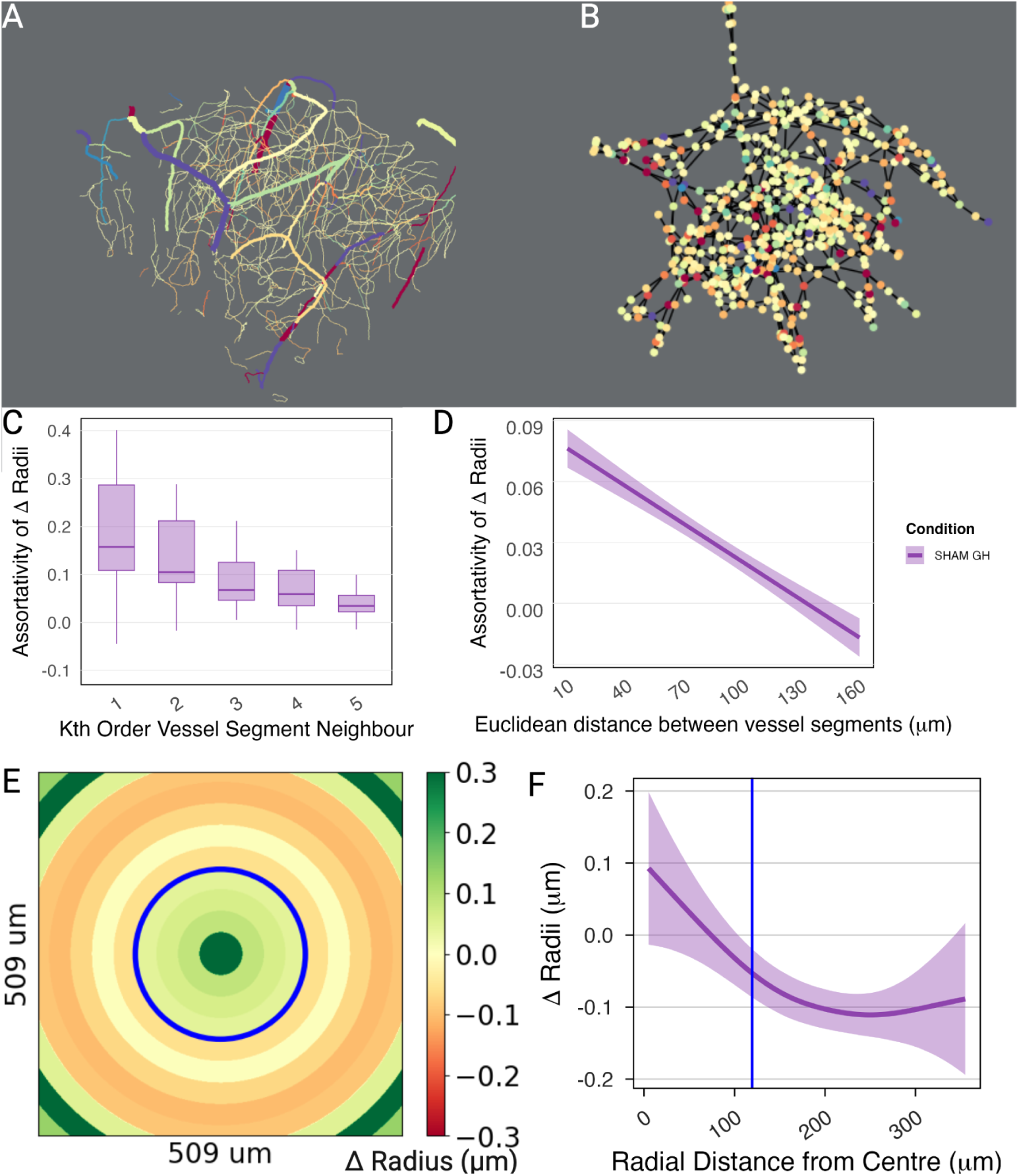
Vascular Network Coordination in SHAM Group Housed mice. **A.** Representative spatial map of radius changes following 4.3 mW/mm^2^ optogenetic stimulation. **B.** Topological network representation of the data in **A**; vessels are depicted as nodes connected to their immediate (1^st^ degree) neighbours via edges. **C.** Assortativity (correlation) of vascular radii changes plotted as a function of distance along the network (degree of separation) between vessel segments. **D.** Spatial assortativity of vascular radii changes between non-adjacent vessels plotted as a function of Euclidean distance. **E.** Average significant vascular radius changes (filtered for absolute changes exceeding 2x baseline SD) aggregated into 30-μm concentric bins radiating from the stimulation centre. **F.** Change in radii as a function of distance from the centre of the photostimulation region of interest (ROI) visualized using ggplot with geom_smooth fitting the significant change in capillary radius as a function of radial distance from the centre of optogenetic stimulation with a GAM. To prevent pseudo-replication from dense vertex-wise data, significant observations were spatially downsampled into 10-μm bins per vessel. The model utilizes cubic regression splines (basis dimension k=5) and a random intercept for subject. Shaded regions represent 95% confidence intervals; non-overlapping bands indicate a robust, statistically significant difference.

### SI Induces Baseline Constrictions Without Altering Reactivity

We next evaluated the effects of chronic social stress on vascular function by comparing SHAM Socially Isolated (SI) mice to the SHAM GH controls (**Figure 4**). At baseline, SI led to significant vasoconstriction. Mean arteriolar radius decreased by 0.83±0.24 μm (p_adj_=0.002) and mean venular radius decreased by 0.96±0.19 μm (p_adj_<1e-4) relative to SHAM GH mice (**Figure 4A**). This structural narrowing was accompanied by a reduction in baseline arteriolar VRBC of 0.70±0.12 mm/s (p_adj_<1e-4) (**Figure 4B**).

**Figure 4:**
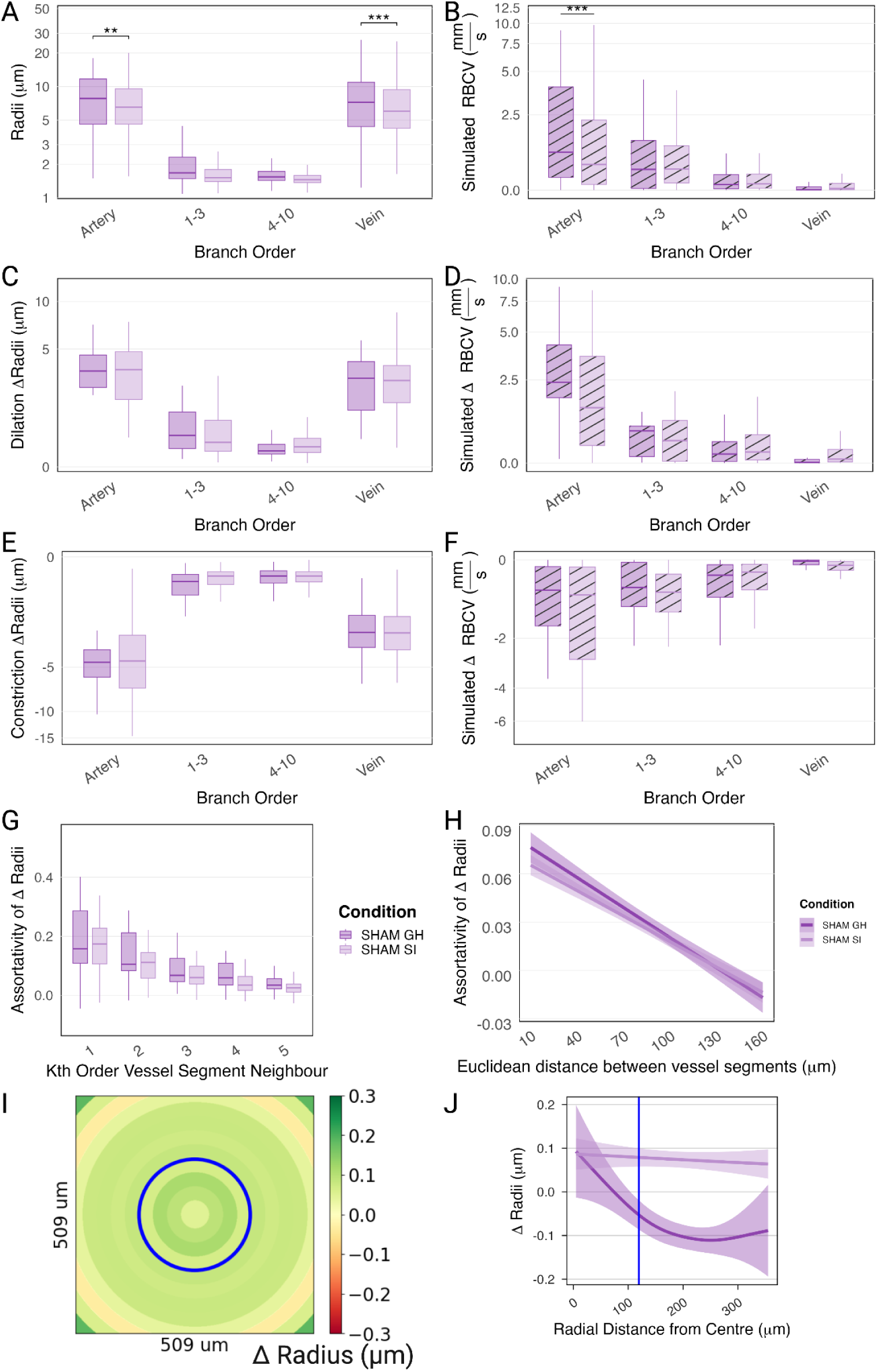
Alterations to Vascular Reactivity Caused by SI. **A.** Baseline vascular radius estimates. Mean arteriolar radius decreased by 0.83±0.24 μm (p_adj_=0.002) and mean venular radius decreased by 0.96±0.19 μm (p_adj_<1e-4) in SHAM SI mice relative to SHAM GH. **B.** Baseline simulated VRBC. Arteriolar velocity decreased by 0.70±0.12mm/s (p_adj_<1e-4) in SHAM SI mice relative to controls. **C–F.** Hemodynamic responses to 4.3 mW/mm^2^ optogenetic stimulation. No significant differences were observed between SHAM SI and SHAM GH mice in dilations (**C**), constrictions (**E**), increases in VRBC (**D**), or decreases in VRBC (**F**). **G.** Topological assortativity of vascular radii changes as a function of distance along the network. **H.** Spatial assortativity of radii changes as a function of Euclidean distance. **I.** Average significant vascular radius changes (filtered for absolute changes >2x baseline SD) aggregated into 30-μm concentric bins radiating from the stimulation centre. **J.** Change in radii as a function of distance from the centre of the photostimulation ROI visualized using ggplot with geom_smooth fitting the significant change in capillary radius as a function of radial distance from the centre of optogenetic stimulation with a GAM, comparing SHAM GH and SHAM SI mice.

Crucially, these changes were limited to vessel calibre. Detailed morphological analysis revealed that vessel length and tortuosity remained unchanged across branch orders (**Figure S3 C-F**). Despite baseline deficits, the hemodynamic reactivity to optogenetic stimulation remained largely intact. We observed no differences between SHAM SI and SHAM GH mice in the magnitude of vessel dilations (**Figure 4C**), constrictions (**Figure 4E**), or evoked changes in VRBC (**Figure 4D, F**) to 4.3 mW/mm^2^ optogenetic stimulation. The lack of significant differences between constrictions, dilations, or VRBC changes in SHAM SI relative to SHAM GH was also observed following 1.1 mW/mm^2^ 458 nm stimulation (**Figure S6**), 4.3 mW/mm^2^ 552 nm photostimulation (**Figure S7**), and forepaw stimulation (**Figure S8**). Furthermore, spatiotemporal analyses, including topological and spatial assortativity **(Figure 4G-H)**, indicated that SI alone did not significantly disrupt neurovascular coupling. To robustly capture spatial dynamics and compare them to healthy controls without artificial precision, vertex-wise data of significant vascular responses (>2x baseline SD) were analyzed using a GAM **(Figure 4I-J)**. While the spatial response profile of SHAM SI mice exhibited central dilations similar to SHAM GH controls, the curves significantly diverged in the periphery. As evidenced by the non-overlapping 95% confidence intervals beginning at 71.5 μm, SHAM SI mice failed to exhibit the peripheral constrictions characteristic of the healthy cohort (up to 359 μm).

### TBI Attenuates Vascular Reactivity and Induces Baseline Constriction

TBI in GH mice produced distinct structural and functional deficits (compared to SHAM controls) (**Figure 5**). Similar to the SI cohort, TBI GH mice exhibited significant baseline vasoconstriction, with arteriolar radii decreasing by 1.16±0.23 μm (p_adj_<1e-4) and venular radii by 1.29±0.19 μm (p_adj_<1e-4, **Figure 5A**). This remodelling was specific to vessel calibre; we observed no significant changes in vessel length or tortuosity across any branch order (**Figure S3 C-F**), indicating that TBI induces vessel lumen narrowing without gross morphological restructuring. Baseline arteriolar VRBC was commensurately reduced by 0.70±0.13 mm/s (p_adj_<1e-4) (**Figure 5B**).

**Figure 5:**
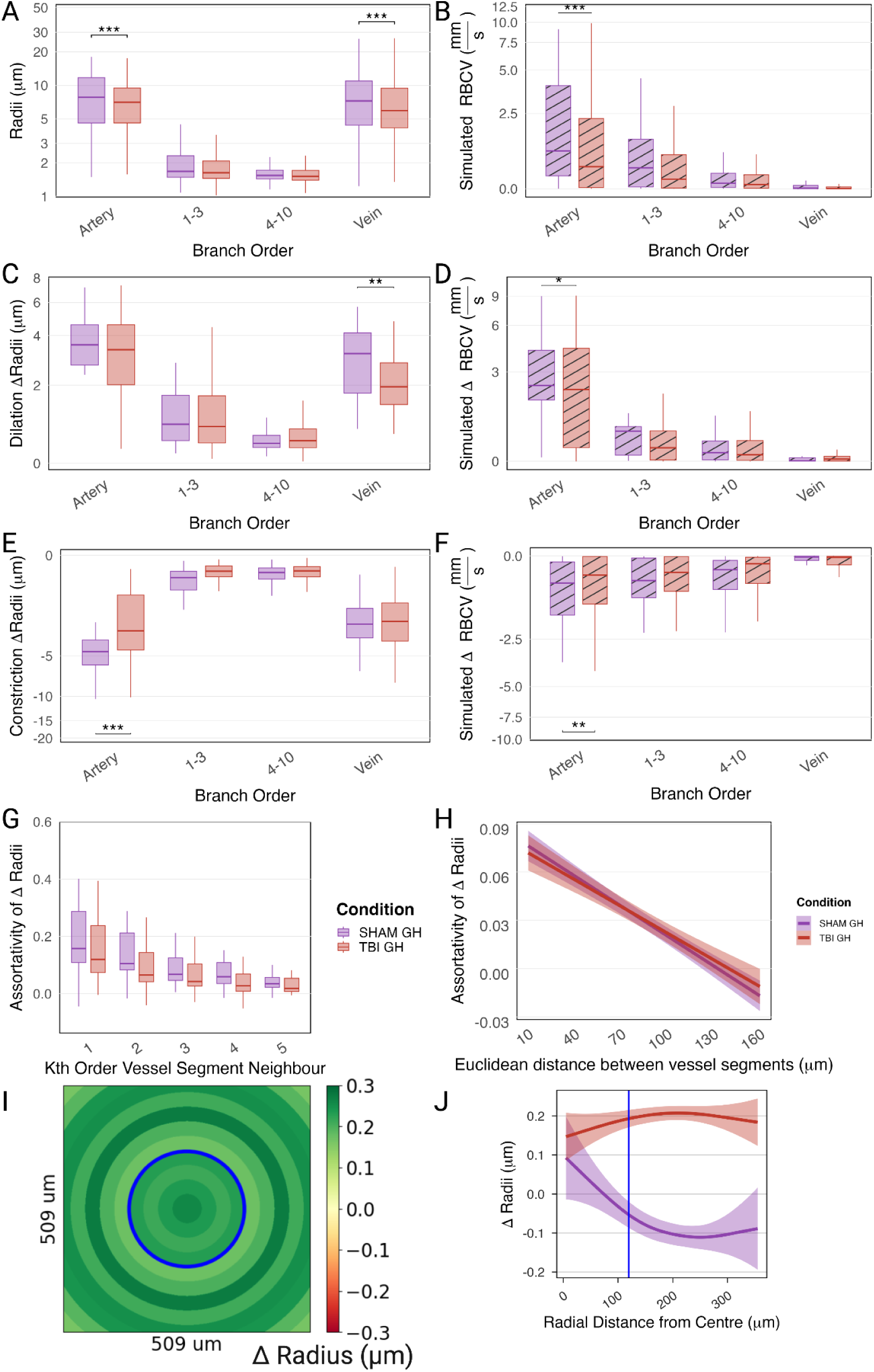
Alterations to Vascular Reactivity Caused by TBI. **A.** Baseline vascular radius comparisons showing a decrease in mean arteriolar (1.16±0.23 μm, p_adj_<1e-4) and venular (1.29±0.19 μm, p_adj_<1e-4) radius in TBI GH mice relative to SHAM GH. **C.** Vascular dilations following 4.3 mW/mm^2^ optogenetic stimulation. TBI GH veins showed a significant decrease in dilation magnitude (0.81±0.23 μm, p_adj_=0.003) **E.** Vascular constrictions following stimulation. TBI GH arteries showed a significant decrease in constriction magnitude (1.76±0.35 μm, p_adj_<1e-4). **B.** Baseline arteriolar velocity was decreased by ( 0.70±0.13 mm/s, p_adj_<1e-4) in TBI GH mice. **D.** Increases in VRBC following stimulation. TBI GH veins showed a significantly blunted VRBC response by (0.55±0.19 mm/s, p_adj_=0.02). **F.** Decreases in VRBC following stimulation. TBI GH arteries showed a significant decrease in magnitude of VRBC decreases by (0.85±0.25 mm/s, p_adj_=0.002). **G–H.** Topological (**G**) and spatial (**H**) assortativity of vascular radii changes were not affected **I.** Average significant vascular radius changes (filtered for absolute changes >2x baseline SD) aggregated into 30-μm concentric bins radiating from the stimulation centre. **J.**Change in radii as a function of distance from the centre of the photostimulation ROI visualized using ggplot with geom_smooth fitting the significant change in capillary radius as a function of radial distance from the centre of optogenetic stimulation with a GAM, comparing SHAM GH and TBI GH mice.

In addition, TBI impaired evoked vascular responses. TBI GH mice displayed a significant decrease in the magnitude of venous dilations (0.81±0.23 μm, p_adj_=0.003, **Figure 5C**) and arteriolar constrictions (1.76±0.35 μm, p_adj_<1e-4, **Figure 5E**). These deficits were present across stimulation intensities; even at lower optogenetic power (1.1 mW/mm^2^ 458 nm), TBI GH arteries exhibited significantly reduced dilation magnitudes (1.05±0.30 μm, p_adj_=0.007) relative to SHAM GH controls (**Figure S6 A**).

These deficits translated into blunted blood flow responses to stimulation: TBI GH veins showed significantly reduced VRBC increases (0.55±0.19 mm/s, p_adj_=0.02, **Figure 5D**), and arteries showed significantly smaller velocity decreases (0.85±0.25 mm/s, p_adj_=0.002, **Figure 5F**). Despite these magnitude changes, network coordination (assortativity) was preserved, with no differences between TBI GH and SHAM GH groups (**Figure 5G-H**).

However, assessing the spatial distribution of these responses revealed injury-induced alterations. To robustly capture spatial dynamics without artificial precision, vertex-wise data of significant vascular responses (>2x baseline SD) were spatially downsampled and analyzed using a GAM **(Figure 5I-J)**. This approach demonstrated that the overall spatial response profile of TBI GH mice significantly differed from SHAM GH controls (p < 0.001). Specifically, as evidenced by the non-overlapping 95% confidence intervals, the response curves significantly diverge beyond 40.4 μm. While SHAM GH controls transition into peripheral constrictions, TBI GH mice exhibit a sustained, abnormal spatial trajectory across the entire remaining radial distance (up to 359 μm), indicating that TBI profoundly alters the healthy spatial gradient of capillary reactivity.

### Combined TBI and SI Severely Blunts Arteriolar Reactivity

Inducing TBI following SI (TBI SI) resulted in the most profound deficits, particularly in arteriolar function and baseline venular structure (**Figure 6**). At baseline, TBI SI mice showed a significant decrease in mean venular radius of 0.82±0.20 μm (p_adj_=0.0003) relative to SHAM GH controls (**Figure 6A**). Whereas TBI GH arteries were constricted relative to SHAM GH, TBI SI arteries displayed a mean *increase* in radius compared to the TBI GH group (0.60±0.22 μm, p=0.005, **Figure S4A**), indicating an interaction between the two stressors.

**Figure 6:**
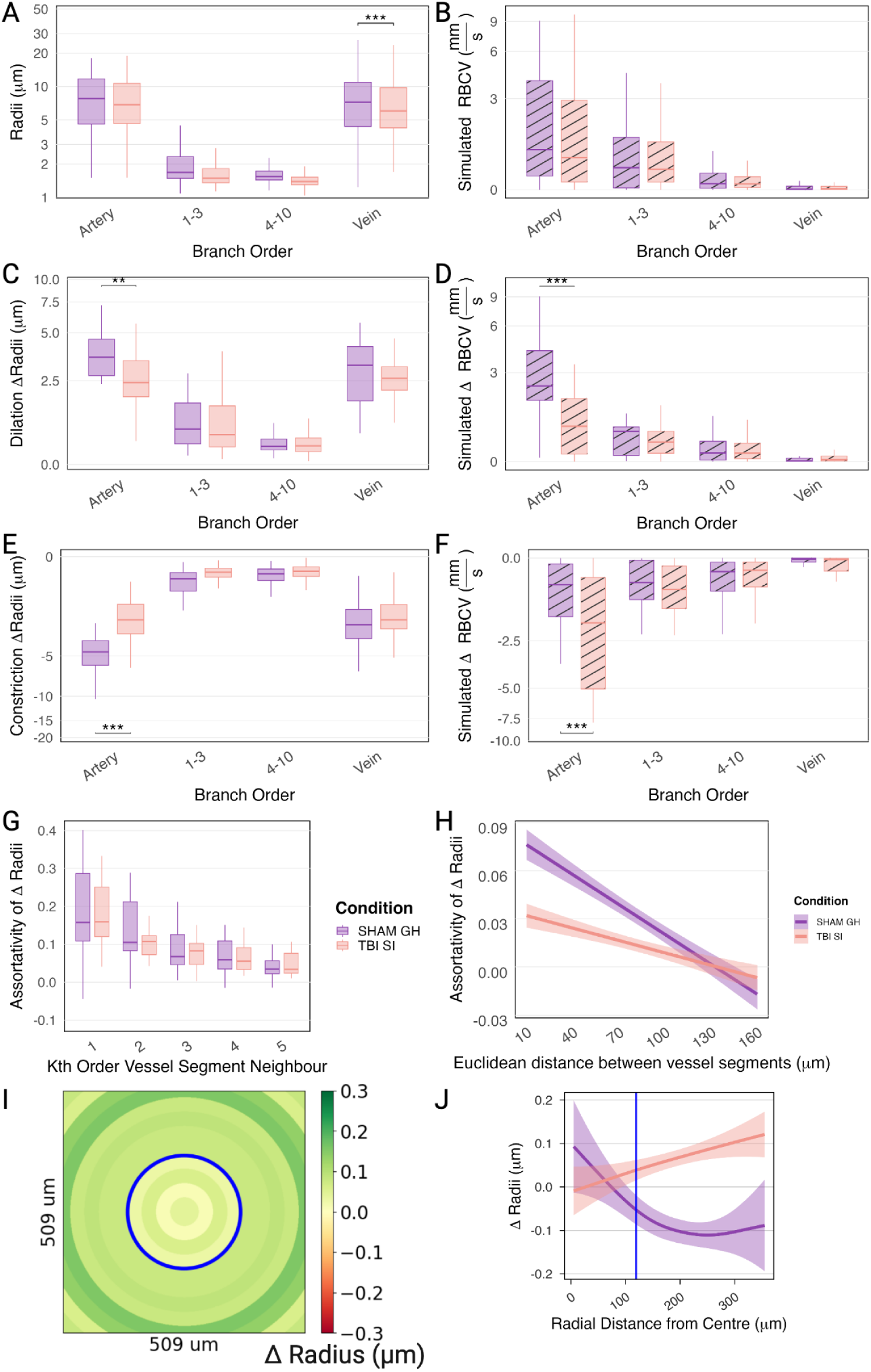
Alterations to Vascular Reactivity Caused by TBI following SI. **A.** Baseline vascular radius comparisons showing a decrease in mean venular radius of 0.82±0.20 μm (p_adj_=0.0003) in TBI SI mice relative to SHAM GH. **C.** Vascular dilations following 4.3 mW/mm^2^ optogenetic stimulation. TBI SI arteries showed significantly blunted dilation magnitudes by (1.07±0.27 μm, p_adj_=0.0008). **E.** Vascular constrictions following stimulation. TBI SI arteries showed significantly decreased constriction magnitudes by (2.40±0.35 μm, p_adj_<1e-4). **B.** Baseline simulated VRBC were indistinguishable. **D.** Increases in VRBC following stimulation. TBI SI arteries showed responses blunted by (2.07±0.23 mm/s, p_adj_<1e-4). **F.** Decreases in VRBC following stimulation. TBI SI arteries showed decreased response magnitudes by (1.23±0.24 mm/s, p_adj_<1e-4). **G.** Topological assortativity of vascular radii changes was unaffected. **H.** Spatial assortativity of vascular radii changes as a function of Euclidean distance was significantly different in TBI SI vs. SHAM GH. **I.** Average significant vascular radius changes (filtered for absolute changes >2x baseline SD) aggregated into 30-μm concentric bins radiating from the stimulation centre. **J.** Change in radii as a function of distance from the centre of the photostimulation ROI visualized using ggplot with geom_smooth fitting the significant change in capillary radius as a function of radial distance from the centre of optogenetic stimulation with a GAM, comparing SHAM GH and TBI SI mice.

Strikingly, the TBI SI cohort exhibited a severe attenuation of arteriolar dilations that was not observed following 4.3 mW/mm^2^ 458 nm optogenetic stimulation in TBI GH or SHAM SI mice. Following stimulation, TBI SI arteries showed significantly blunted dilation magnitudes (by 1.07±0.27 μm, p_adj_=0.0008, **Figure 6C**) and significantly decreased constriction magnitudes (by 2.40±0.35 μm, p_adj_<1e-4, **Figure 6E**). This attenuation was supralinear; TBI SI arteriolar responses were significantly reduced beyond combined changes in the TBI GH and SHAM SI cohorts (**Figure S4 B, E**), and these deficits persisted even under lower-power stimulation (1.1 mW/mm^2^, **Figure S6**).

Consistent with the notion of arterioles being key for healthy cortical perfusion^45,46^, this loss of arteriolar reactivity had major functional consequences for blood flow regulation. TBI SI arteries showed a significantly blunted capacity to increase VRBC (2.07±0.23 mm/s, p_adj_<1e-4, **Figure 6D**) or decrease it (1.23±0.24 mm/s, p_adj_<1e-4, **Figure 6F**).

To robustly capture spatial dynamics without artificial precision, vertex-wise data of significant vascular responses (>2x baseline SD) were spatially downsampled and analyzed using a GAM **(Figure 6I-J)**. This approach demonstrated that the overall spatial response profile of TBI SI mice significantly differed from SHAM GH controls (p = 0.001). Specifically, the response curves significantly diverge between approximately 102.5 μm and 355 μm from the stimulation centre. While SHAM GH controls maintain a spatial profile that begins with a central dilation and gradually transitions into a true peripheral constriction (dropping below baseline), TBI SI mice exhibit a blunted central dilation and completely fail to constrict in the periphery. Instead, the TBI SI network maintains a low-level, sustained dilation across the entire distal region. This distinct spatial profile indicates that the combination of TBI and SI drives a unique neurovascular dysfunction characterized by a flattened, globally attenuated reactivity profile that lacks both robust central hyperaemia and healthy peripheral constriction.

## Discussion

Our data demonstrate that TBI, when compounded with chronic stress induced via SI, fundamentally disrupts the long range coordination of the capillary network and severely blunts arteriolar reactivity. While stress alone induced baseline vasoconstriction without impairing functional reactivity, and TBI alone attenuated evoked responses, the combination of TBI and SI produced the most profound functional deficits. Specifically, TBI+SI dismantled the network’s ability to smoothly distribute blood flow, flattening the healthy spatial gradient by exhibiting severely blunted central dilations and failing to mount healthy peripheral constrictions.

Reduced neurovascular coupling and hypoperfusion in chronic stages of TBI are well documented and correlate with adverse outcomes ^22,23,47–50^. Uncovering the underlying microvascular mechanisms that sustain these deficits is critical, as chronic hypoperfusion can lead to local hypoxia and sustain secondary injury ^16,24,26,51^. Neurovascular coupling deficits in TBI, i.e. attenuation of blood flow increases following focal neuronal activation, have been widely reported on in macroscopic cerebral blood flow (CBF) and microscopic VRBC-focused studies. Specifically, the diminished arteriolar constrictions and dilations we observed directly align with the decreased vascular reactivity reported in MRI studies ^22,52,53^. This is complemented by simulations showing decreased ΔVRBC reactivity, which aligns with Adams et al.’s finding that cerebrovascular reactivity and CBF were reduced and Van Hameren et al.’s demonstration of impaired blood flow during at least part of the vascular response phase, and with our earlier study showing reduced VRBC in arteries^22,54^. The most dramatic contrasts for the 1.1 mW/mm^2^ and 4.3 mW/mm^2^ 458 nm optogenetic stimulations were observed in arteries and veins, where TBI resulted in significant decreases in arteriolar reactivity and venous dilations, and TBISI led to the greatest reduction in arteriolar reactivity. SI-induced chronic stress perpetuates these deficits in arteriolar reactivity, as chronic stress can lead to increased corticosterone, which suppresses vasodilatory responses in vascular smooth muscle cells ^55,56^. Our simulations also indicated the greatest reduction in the ability of vascular networks to increase arterial blood flow in the TBI+SI cohort. This severe impairment aligns with, and demonstrates a worsening of, the reduced blood flow modulation previously reported in TBI alone ^22^.

A major novelty of this work lies in elucidating the network-level mechanisms underlying these deficits. Functionally, endothelial cell-mediated propagation of electrical currents unifies vascular responses across adjoining vessels, whereas long-distance mechanisms can either lead to antagonistic or synergistic responses. By employing graph-based assortativity analysis, we distinguished between these two modes of coordination. There were no cohort-wise differences in assortativity between directly connected vessels, implying that gap junctions between endothelial cells remain intact and propagated K+ currents were unaffected ^57^. In contrast, TBI+SI significantly attenuated assortativity in distant, non-directly connected vessels. In all groups except those with TBI SI, the assortativity of radii decreased as the distance between linked vessels increased. The ability of vessels to coordinate local increases in vascular radii may thus be impaired in TBISI, potentially reducing the vasculature’s ability to distribute blood flow. Mechanisms that act on distant vessels that are not immediately connected, such as astrocyte-mediated NO signalling and interpericyte tunnelling nanotubules, are likely impaired in TBI ^58–60^.

In healthy mice, vessels dilated focally in the region of neuronal activation while surrounding vessels constricted, redistributing blood towards areas with higher neuronal activity ^61,62^. Although the redistribution of blood flow to areas of increased neuronal activity has been established for a long time ^45^, how these opposing changes in vascular tone are spatially coordinated across the broader capillary network remains unclear.

Following TBI and/or SI, this healthy spatial gradient was disrupted via distinct pathological trajectories. SHAM SI and TBI GH mice failed to transition into peripheral constrictions, maintaining abnormally sustained dilated profiles. Conversely, TBI SI mice exhibited a globally attenuated response, failing to mount robust central dilations and similarly failing to constrict in the periphery. In all cases, the physiological contrast between the activation core and its surrounding area was severely diminished. This finding is in agreement with assortativity patterns seen in TBI SI mice, where no distance-dependent vascular responses were observed, unlike SHAM GH mice, in which nearby vessels responded similarly, while distant vessels responded differently. Since SHAM SI and TBI GH showed intermediate patterns, TBI and SI may have synergistic effects on vascular coordination and spatial organization. Moreover, reduced spatial assortativity was associated with impaired redistribution of blood flow increases and decreases relative to the neuronal activation core, suggesting that this breakdown in network-level spatial organization fundamentally underpins the functional failure to redistribute blood flow.

### Limitations

Several limitations are noteworthy. We employed optogenetic stimulation to activate pyramidal neurons in the cortex, thereby eliciting long-lasting (at least 45 seconds) changes to VRBC ^22,63,64^, with recent work showing vascular tone modulation lasting for minutes when examining diameters ^65^. This targeted neuronal stimulation contrasts with widefield optogenetic stimulations of neurons with fibre optic probes flashing blue light over large areas which leads to shorter-lasting vascular responses (under 20 seconds)^64^. We utilized the resonant scanning technique and maximized our imaging volumes to capture as many vessels as possible in under 45 seconds, thereby maximizing our imaging field of view (FOV) while compromising temporal resolution. While our volumetric scanning approach limited temporal resolution, preventing analysis of kinetics, it enabled an unprecedented, network-wide view of hundreds of vessels simultaneously. This coverage was essential for elucidating deficits in spatial coordination, a significant advancement over traditional line-scanning techniques that can interrogate only a handful of vessels at once. While we observed similar trends in reduced arteriolar reactivity in the forepaw stimulation data, the results did not reach significance, in part due to the low number of subjects who underwent these stimulations and scans (n=24, TBI SI=1). Additionally, our blood flow (VRBC) estimates are computationally constrained by the limited imaging FOV, which lacks data on broader upstream and downstream vascular resistance.

### Targeting Microvascular Coordination in TBI

Cerebrovascular impairments have been suggested as a biomarker for individuals who suffer from adverse outcomes post TBI. Long-term inability to modulate blood flow is thought to lead to the development of chronic pockets of hypoxia in the tissue, limiting the recovery^18^. Here, we demonstrated that TBI reduces arteriolar reactivity, but critically, it is the compounding effect of chronic stress (TBI+SI) that drives the severe reduction in long-distance coordination of vascular changes throughout the capillary bed. These results identify potential microvascular targets (e.g. buffering corticosterone-induced suppression of arteriolar smooth muscle cells, and restoring distant capillary coordination via astrocytic NO signalling or interpericyte tunnelling nanotubes) for further study in the development of therapeutics that could restore cerebrovascular function. Furthermore, our findings indicate that chronic stress increases the brain’s susceptibility to lasting injury from TBI, with exacerbated cerebrovascular dysfunction. Because the network-wide coordination of capillary blood flow is actively driven by astrocytes and pericytes, targeting astrocyte and pericyte function represents a promising avenue for restoring blood flow regulation in TBI patients, particularly those with comorbid stress-related conditions.

### Conclusion

We demonstrated that TBI disrupts neurovascular coupling within capillary networks by reducing arteriolar reactivity and impairing capillary coordination. TBI and SI resulted in blunted arteriolar reactivity, suggesting the metabolic demand during neuronal activation may not be met effectively. Our simulation predicting blood flow through the observed network revealed a diminished capacity to enhance blood flow to areas of increased neuronal activity in TBI SI mice. While coordination between directly connected vessels remained intact, coordination over long distances was diminished. This finding indicates that long-range mechanisms of vascular coordination are compromised during the chronic stages of TBI comorbid with SI. Alterations in spatial coordination were accompanied by an impaired ability to properly graduate blood flow across the network; individual stressors resulted in an abnormal failure to constrict the periphery, while compounded stressors (TBI+SI) drove a flattened, globally attenuated response lacking robust central hyperaemia. These findings highlight potential targets for future treatments, particularly focusing on restoring arteriolar smooth muscle cell reactivity and targeting astrocyte or pericyte pathways to rescue distant capillary coordination. These targets parallel clinical realities, where preexisting vulnerabilities deeply influence three-month patient outcomes ^66^. Addressing these stress-exacerbated microvascular failures provides a critical translational link to mitigating the sustained, real-world post-concussive deficits observed in the general population. Additionally, we advanced methodological approaches by deploying the NOVAS3D volumetric scanning pipeline to extract full microvascular geometries and classify distinct vessel types. This comprehensive structural mapping directly enabled our network-wide blood flow simulations and graph-based analysis of capillary coordination. By elucidating the specific network-level mechanisms underlying neurovascular dysfunction, this work provides a mechanistic foundation for understanding sustained deficits post TBI and identifies potential targets for intervention to improve long-term outcomes.

## Novelty and Significance

### What Is Known?

● Chronic stress strongly predicts poor recovery and permanent disability following traumatic brain injury (TBI).
● Mechanisms underlying post-concussive cerebrovascular dysfunction, hypoperfusion, and secondary disease progression remain unclear.

### What New Information Does This Article Contribute?

● Comorbid TBI and chronic stress blunts arteriolar reactivity, reducing vascular dilation magnitudes by about 30% and constriction magnitudes by half following neuronal activation.
● Combined injury disrupts long-range functional capillary coordination, despite intact local vessel coordination.
● Impaired spatial coordination of cerebrovascular calibre changes blurs blood flow redistribution, diminishing epicentral dilations and peripheral constrictions.

### Summary

Understanding sustained impairment of microvascular coordination is critical for improving neurophysiological recovery after TBI. Using our NOVAS3D analysis pipeline, we evaluated network-level neurovascular coupling in mice with TBI and comorbid chronic stress. Although vascular topology remains structurally intact, combined injury abates long-range capillary synchronization. This uncoupling is driven by severe attenuation of arteriolar reactivity, yielding an estimated 70% reduction in red blood cell velocity responses to neuronal activation. Consequently, the network loses its capacity to spatially redistribute blood flow and exhibits an attenuated reactivity profile. These findings establish that altered spatial coordination and blunted reactivity underpin chronic post-TBI hypoperfusion, identifying them as critical therapeutic targets.

## Acknowledgments

We thank Calcul Québec and the Digital Research Alliance of Canada for compute resources. Supported by Canadian Institutes of Health Research grants PJT376309, PJT156179, and PJT178059. M.W.R. was supported by QEII/Sunnybrook and Women’s College Health Sciences Centre GSST and VAST funding. B.S. and M.G. are supported by the Canada Research Chair program (CRC-2018-00042, CRC-2021-00374).

## Author contributions

MWR, AD, SP, and JRM conducted data acquisition. MWR, AD, SP, AA, YD, and JO performed data analysis. MK, AD, SP, and MH performed surgical and experimental preparation. MWR, JRM, MB, CH, JGS, MG, and BS designed the experiment. All authors contributed to manuscript preparation and approved the final submission.

## Code and data availability

The NOVAS3D pipeline used for data analysis is available here: https://github.com/AICONSlab/novas3d. Datasets will be made public and uploaded to the FRDR.

## Abbreviations

2PFM: Two-photon fluorescence microscopy
BPM: Beats per minute / Breaths per minute
CBF: Cerebral blood flow
ChR2: Channelrhodopsin-2
FOV: Field of view
GAM: Generalized Additive Model
GH: Group Housed
NOVAS3D: Network of Vascular Analysis 3D pipeline
RBC: Red blood cell
ROI: Region of interest
SD: Standard deviation
SI: Social isolation
SNR: Signal-to-noise ratios
TBI: Traumatic brain injury
THY1: THY1-ChR2-EYFP (mouse line)
VRBC: Red blood cell velocity.

